# cGAS-STING is responsible for aging of telomerase deficient zebrafish

**DOI:** 10.1101/2024.03.11.584360

**Authors:** Naz Şerifoğlu, Giulia Allavena, Bruno Bastos-Lopes, Marta Marzullo, Pavlos Bousounis, Eirini Trompouki, Miguel Godinho Ferreira

**Affiliations:** Institute for Research on Cancer and Aging of Nice (IRCAN), CNRS UMR7284, INSERM U1081, Université Cote d’Azur, 06107 Nice, France; Instituto Gulbenkian de Ciência, Oeiras, Portugal; Department of Cellular and Molecular Immunology, Max Planck Institute of Immunobiology and Epigenetics, Freiburg, Germany; Faculty of Biology, University of Freiburg, Freiburg, Germany

**Keywords:** Telomerase, cGAS-STING, inflammation, aging, zebrafish

## Abstract

Telomere shortening occurs in multiple tissues throughout aging. When telomeres become critically short, they trigger DNA damage responses and p53 stabilization, leading to apoptosis or replicative senescence. *In vitro*, cells with short telomeres activate the cGAS-STING innate immune pathway resulting in type I interferon inflammation and senescence. However, the consequences of these events to the organism are not yet understood. Here, we show that *sting* is responsible for premature aging of telomerase-deficient zebrafish. We generated *sting-/- tert-/-* double mutants and observed a thorough rescue of *tert-/-* phenotypes. At the cellular level, lack of cGAS-STING in *ter t* mutants resulted in reduced senescence, increased cell proliferation, and low inflammation despite similar short telomeres. Critically, absence of *sting* function resulted in dampening of the DNA damage response and low p53 levels. At the organism level, *sting-/- tert-/-* zebrafish regained fertility, delayed cachexia, and cancer incidence, resulting in increased healthspan and lifespan of telomerase mutants.

## Introduction

Telomerase deficiency in humans results in the development of telomere biology disorders (TBDs) that include idiopathic pulmonary fibrosis, dyskeratosis congenita and aplastic anemia^1^. Common aspects of TBDs relate to accelerated telomere shortening, loss of tissue regeneration, premature aging phenotypes and shorter life span^2,3^. Zebrafish possess human-like telomere lengths that shorten to critical lengths during their lifetime. Like human TBDs, telomerase-deficient zebrafish (*tert-/-*) have accelerated telomere shortening, low cell proliferation, tissue damage, and reduced lifespan ^4–6^. *tert-/-* zebrafish also develop chronic inflammation, increased infections, and accelerated incidence of cancer^7,8^. Like in humans^9^, we have recently shown that not all organs age at the same rate^8^. The zebrafish intestine becomes dysfunctional earlier and triggers systemic aging. Reduced proliferation of intestine cells results in loss of tissue integrity, microbiota dysbiosis and systemic inflammation^8^.

Type I interferon response and secretion of pro-inflammatory cytokines through activation of cGAS-STING is triggered by DNA damage, including telomeric damage, leading to the formation of micronuclei (MN)^10–13^. The cytosolic DNA sensor, cGAS (cGMP-AMP synthase), becomes activated upon binding to double-stranded DNA, encompassing both microbial and self-DNA^14^. Upon recognition of cytosolic DNA, cGAS initiates the production of the second messenger cGAMP that binds and activates the adaptor protein STING^15,16^. Subsequently, STING recruits TBK1 (TANK-binding kinase 1) triggering the activation of IRF3 (IFN regulatory factor 3). This activation cascade leads to the generation of type I interferons and inflammatory cytokines^14^. cGAS-STING is involved in DNA repair, DNA damage responses (DDR) and cell senescence (reviewed in^17^). These effects are achieved through the expression of interferon-stimulated genes (ISGs) and the senescence-associated secretory phenotype (SASP)^12,16^. Secretion of these molecules modulates the proliferative capacity of surrounding cells in a paracrine manner, propagating the senescence status (paracrine-SASP)^18^.

Type I interferon response is increasingly linked to aging and neurodegenerative diseases across different species. A recent significant study by the Ablasser lab using naturally aged mice revealed that cGAS–STING signaling plays a pivotal role in the age-related type I interferon response in neurodegeneration^19^. Transcriptional profiling revealed that cGAS–STING activation initiated a gene expression program shared between neurodegenerative diseases and natural aging. The authors also observed that there was an accumulation of mitochondrial DNA in the cell cytoplasm of microglial cells, providing a potential mechanism through which the cGAS-STING pathway may contribute to inflammation in the aging brain^19^.

*In vitro* studies using human primary cells recently showed that cGAS-STING is activated by short or dysfunctional telomeres (reviewed in^13^). However, it is currently unknown what are the consequences of activation of cGAS-STING in response to short telomeres at the organism level. Here, we show that telomere shortening triggers cGAS-STING pathway in skin, testis, kidney marrow (the adult hematopoietic organ in fish) and intestine of zebrafish. This results in type I interferon response, elevation in senescence levels and reduction in proliferative capacity of these tissues. Absence of cGAS-STING restores cell proliferation, suppresses accelerated aging and increases in both health and lifespan of the telomerase deficient zebrafish.

## Results

### Telomere shortening activates the cGAS STING and type I interferon *in vivo*

Our previous studies show that telomerase deficient zebrafish undergo accelerated systemic inflammation^5–8,20,21^. Analysis of gene expression of proliferative (gut) and non-proliferative tissues (muscle) derived from aged *tert-/-* (9-months old) and WT (36-months old) zebrafish highlighted several genes related to type I interferon response^20^. To investigate if short telomeres trigger type I inflammation and accelerated aging through the cGAS-STING pathway, we combined *sting* zebrafish loss of function mutants (*sting*^sa35634/sa35634^, hereby referred to as *sting-/-*) with telomerase deficient zebrafish (*tert*^hu3430/hu3430^ or *tert-/-*) extensively characterized in our previous studies^5–8,21^. Upon incross of double heterozygous fish, we first investigated if *tert-/- sting-/-* double mutants had short telomeres comparable to their *tert- /*si*-*blings. We observed that, by 9 months of age, *tert-/- sting-/-* had similar mean telomere length to *tert-/-* single mutants in the skin and intestine (Figure 1A-D, Supplementary figure 1A-D). In testis, *tert-/- sting-/-* zebrafish had slightly shorter telomeres than *tert-/-* (Figure 1B, Supplementary Figure 1B) and slightly longer telomeres in the kidney marrow (Figure 1C, Supplementary Figure 1C). Overall, not only telomere length was similar between *tert-/-* and *tert-/- sting-/-*but were significantly shorter than WT and *sting-/-* siblings.

**Figure 1:**
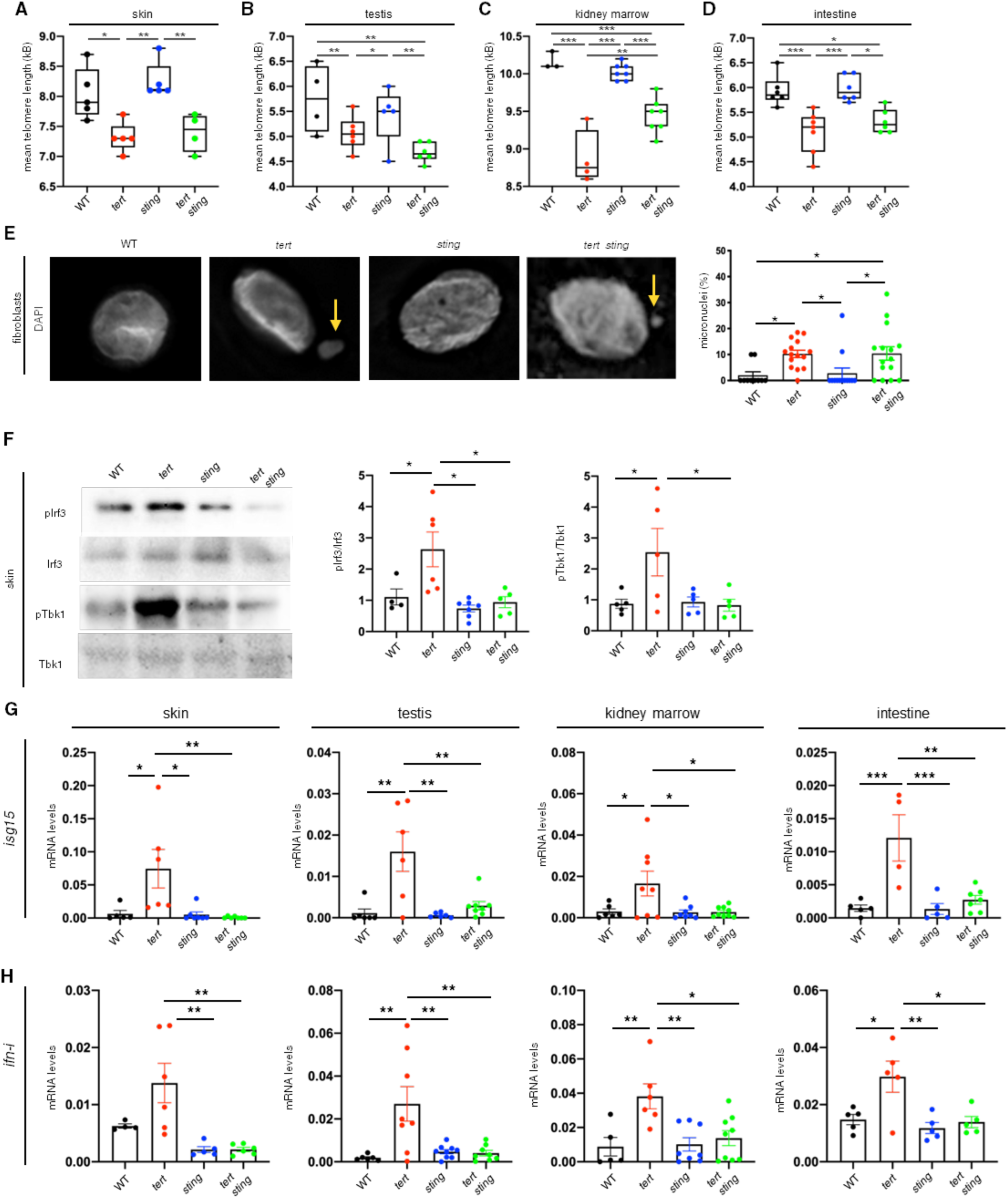
Telomere shortening activates cGAS-STING pathway. **a**, quantification of mean telomere length measured by TRF analysis in the skin (n_WT_= 5, n*_tert-/-_*=5, n*_sting-/-_*=5, *_tert-/-sting-/-_* =4, WT *vs tert-/-* p= 0.017,tert-/- *vs sting-/-* p= 0.002, *sting-/- vs tert-/- sti n*p*g*=0*- /*.0*-*08). **b**, quantification of mean telomere length measured by TRF analysis in the testis (n_WT_= 4, n*_tert-/-_*=6, n*_sting-/-_*=5, *_tert-/-sting-/-_* =6, WT *vs tert-/-* p=0.001, WT *vs tert-/- sting-/-* p<0.001, *tert-/- vs sting-/-* p= 0.016, *sting-/- vs tert-/- sting- /*p*-*=0.001). **c**, quantification of mean telomere length measured by TRF analysis in the kidney marrow (n_WT_= 3, n*_tert-/-_*=4, n*_sting-/-_*=7, *_tert-/-sting-/-_*=7, WT *vs tert-/-* p< 0.001, WT *vs tert-/- sting* p*-*=*/-* 0.001, *tert-/- vs sting-/-* p< 0.001, *sting-/- vs tert-/- sting-/-* p<0.001, *tert-/- vs tert-/- sting-/-* p=0.003). **d**, quantification of mean telomere length measured by TRF analysis in the intestine (n_WT_= 6, n*_tert-/-_*=6, n*_sting-/-_*=6, *_tert-/-sting-/-_* =6, WT vs *tert-/-* p= 0.001, WT vs *tert-/- sting-/-* p=0.015, *tert-/- vs sting-/-* p<0.001, *sting-/- vs tert-/- sting-/-* p=0.008, p= 0.623). **e**, representative immunofluorescence images and quantifications of MN formation in the fibroblasts derived from skin (n_WT_= 1, n*_tert-/-_*=1, n*_sting-/-_*=1, *_tert-/-sting-/-_*=1, WT vs *tert-/-* p= 0.040, WT vs *tert-/- sting-/-* p=0.034*, tert-/-* vs *sting-/-* p= 0.047, *sting-/-* vs *tert-/- sting-/-* p=0.040). **f**, representative western blot images and quantification of downstream targets of cGAS-STING pathway (n_WT_= 4-5, n*_tert-/-_*=5-6, n*_sting-/-_*=5-7, *_tert-/-sting-/-_*=5, p-Irf3/Irf3: WT vs *tert-/- p= 0.049, tert-/- vs tert-/- sting-/- p= 0.011; p-Tbk1/Tbk1: WT vs tert-/- p= 0.035, tert-/- vs sting-/- p= 0.002, tert-/- vs tert-/- sting-/- p= 0.042*). **g**, RT-qPCR analysis of *isg15* gene expression in the skin, testis, kidney marrow and intestine (n_WT_= 5-6, n*_tert-/-_*=4-8, n*_sting-/-_*=5-8, *_tert-/-sting-/-_*=7-9, skin: WT vs *tert-/- p= 0.024, sting-/- vs tert-/- p=0.011, N=5-7, tert-/- vs tert-/- sting-/- p=0.007; testis: WT vs tert-/- p=0.001, sting-/- vs tert-/- p=0.001, tert-/- vs tert-/- sting-/- p=0.002; kidney marrow: N=6-9, WT vs tert-/- p=0.045, sting-/- vs tert-/- p=0.022, tert-/- vs tert-/- sting-/- p= 0.019; intestine: WT vs tert-/- p<0.001, sting-/- vs tert-/- p<0.001, tert-/- vs tert-/- sting-/- p=0.001*). **h**, RT-qPCR analysis of *ifn-i* gene expression in the skin, testis, kidney marrow and intestine (n_WT_= 4-6, n*_tert-/-_*=5-8, n*_sting-/-_*=5-8, *_tert-/-sting-/-_*=6-9, skin: *sting-/- vs tert-/- p=0.004*, *tert-/-* vs *tert-/- sting-/- p=0.002; testis: WT vs tert-/- p=0.003, sting-/- vs tert-/- p=0.003, tert-/- vs tert-/- sting-/- p=0.003; kidney marrow: WT vs tert-/- p=0.008, sting-/- vs tert-/- p=0.004, tert-/- vs tert-/- sting-/- p= 0.011; intestine: WT vs tert-/- p=0.020, sting-/- vs tert-/- p=0.005, tert-/- vs tert-/- sting-/- p=0.014*.) Data are presented as the mean ± s.e.m.; *p<0.05; **p<0.01, ***p<0.001, using a one-way ANOVA and post hoc Tukey test.

Previous studies using cell lines reported that short and dysfunctional telomeres lead to the formation of MN^10–12^. To test whether we would observe an increase of MN upon telomere shortening in zebrafish, we derived fibroblasts from the skin of 9-months old animals. We observed a ten-fold increase in MN formation in *tert-/-* zebrafish when compared to the WT and *sting-/-* siblings. Strikingly, *tert-/- sting-/-* had similar levels of MN as *tert-/-* siblings (Figure 1E). Thus, telomere shortening results in MN accumulation *in vivo*.

After confirming the presence of shorter telomeres and MN in *tert-/-* and *tert-/- sting-/-* zebrafish, we investigated if the cGAS-STING pathway was active *in vivo* by quantifying its downstream targets. Therefore, using skin of 9-months-old zebrafish, we analyzed the phosphorylation status of zebrafish Tbk1 and Irf3 and the transcription level of two members of type I interferon response, *isg15* and *ifn-i* (Figure 1F-H). Comparing *tert-/-* to WT zebrafish we observed ca. 3-fold increase in p-Irf3 and 2-fold increase in p-Tbk1 (Figure 1F). Consistently, expression of *isg15* and *ifn-i* was increased by 10- and 2.5-fold, respectively (Figure 1G-H). However, the phosphorylation profile of Tbk1 and Irf3 in *tert*-/- *sting-/-* mutants was similar to their WT and *sting-/-* siblings (Figure 1F). *tert*-/- *sting-/-* mutants also lacked type I interferon response, as observed by the reduced levels of *isg15* and *ifn-i* (Figure 1G-H).

We expanded our study to other proliferative tissues like testis, kidney marrow and intestine since they all show signs of inflammation in aging zebrafish^5,8,21^. Comparing transcription levels of *tert-/-* fish to WT fish, we observed 15-fold increase in expression for *isg15* and *ifn-i*in testis; 20-fold increase in *isg15 and* 5 fold increase in kidney marrow; and, lastly, 7.5-fold increase in *isg15* transcript and 2.5-fold increase in *ifn-i* in the intestine. Importantly, inactive cGAS-STING pathway rescued the *isg15* and *ifn-i* levels of *tert-/- sting-/-* to those observed in WT zebrafish in all tissues analyzed (Figure 1G, H). These results show that, despite the presence of shorter telomeres and MN, the cGAS-STING pathway is inactive in *tert*-/- *sting-/-* mutants.

Type I interferon response can be initiated by mobilization of transposable elements (TEs). During replicative senescence of human fibroblasts, L1 retrotransposable elements become transcriptionally derepressed and activate a type-I interferon response^22^. TE transcription leads to robust activation of RIG-I and MDA5 that recognize dsRNA and ssRNA and initiate a signaling cascade, involving MAVS oligomerization, IRF3/IRF7 activation and the expression of type I interferon response. We looked for the deregulation of TEs expression in zebrafish testis, kidney marrow and intestine (Supplementary Fig. 2A-B). Multidimensional scaling (MDS) plots of TE expression were non-overlapping for WT and *tert*-/- mutant in all tissues analyzed (Supplementary Fig. 2A). Telomerase-deficient fish showed a clear upregulation of TEs in testis and kidney marrow and, to a lesser extent, in the gut.

We then investigated whether the expressions of the RNA sensors *rig-I*, *mda5* and *mav s* were increased in telomerase-deficient animals. Similar to TE expression, *rig-I* and *mda5* were overexpressed in *tert-/-* kidney marrow and testis, respectively, compared to WT animals, but not the intestine (Supplementary Fig. 2C-D). Recently, the Karlseder lab showed that both DNA sensing (cGAS-STING) and RNA sensing (ZBP1-MAVS) innate immunity pathways orchestrate telomere replicative crisis (M2) ^23^. Their model proposes that short telomeres and MN trigger cGAS-STING, whereas ZBP1 senses the telomeric lncRNA TERRA and activates MAVS. Since both pathways result in the activation of type I interferon response, we tested if ZBP1-MAVS could also contribute to IFN expression. As zebrafish lack a clear *zbp1* gene orthologue, we relied solely on expression of *mavs* for this analysis. *mavs* mRNA was significantly increased in the testis, but not in kidney marrow and intestine (Supplementary Figure 2E). Thus, we observed that telomere shortening triggers multiple responses that include TE expression deregulation and RNA sensor pathway activation which, in turn, reinforce the expression of type I interferon.

### cGAS-STING is required for increased p53 levels in presence of DNA damage

We previously reported that telomere shortening leads to increased p53 levels as a consequence of activation of DDR in zebrafish ^5,6,8,21^. To identify the cause for activation of cGAS-STING in prematurely aged *tert*-/- mutants, we analyzed DNA damage and activation of DDR in 9-month-old animals. First, we quantified the number of phosphorylated H2AX (γ-H2AX) stained cells by immunofluorescence in the tissues of interest. As in our previous work^8^, γ-H2AX staining was dispersed through the nucleoplasm and aggregated in foci (Figure 2A insets). *tert-/-* zebrafish displayed 3-fold increase of γ-H2AX stained cells in the skin (Figure 2B), 4-fold in the testis (Figure 2C) and 5-fold increase both in the kidney marrow (Figure 2D) and the intestine (Figure 2E) when compared to WT and *sting-/-* mutants. Contrarily, *tert-/- sting-/-* mutants showed similar levels of γ-H2AX compared to the *tert-/-* sibling. (Figure 2A-D). Thus, consistent with shorter telomere length in *tert-/-* and *tert-/- sting-/-* mutants, we observed similar levels of the DNA damage marker γ-H2AX in these tissues.

**Figure 2:**
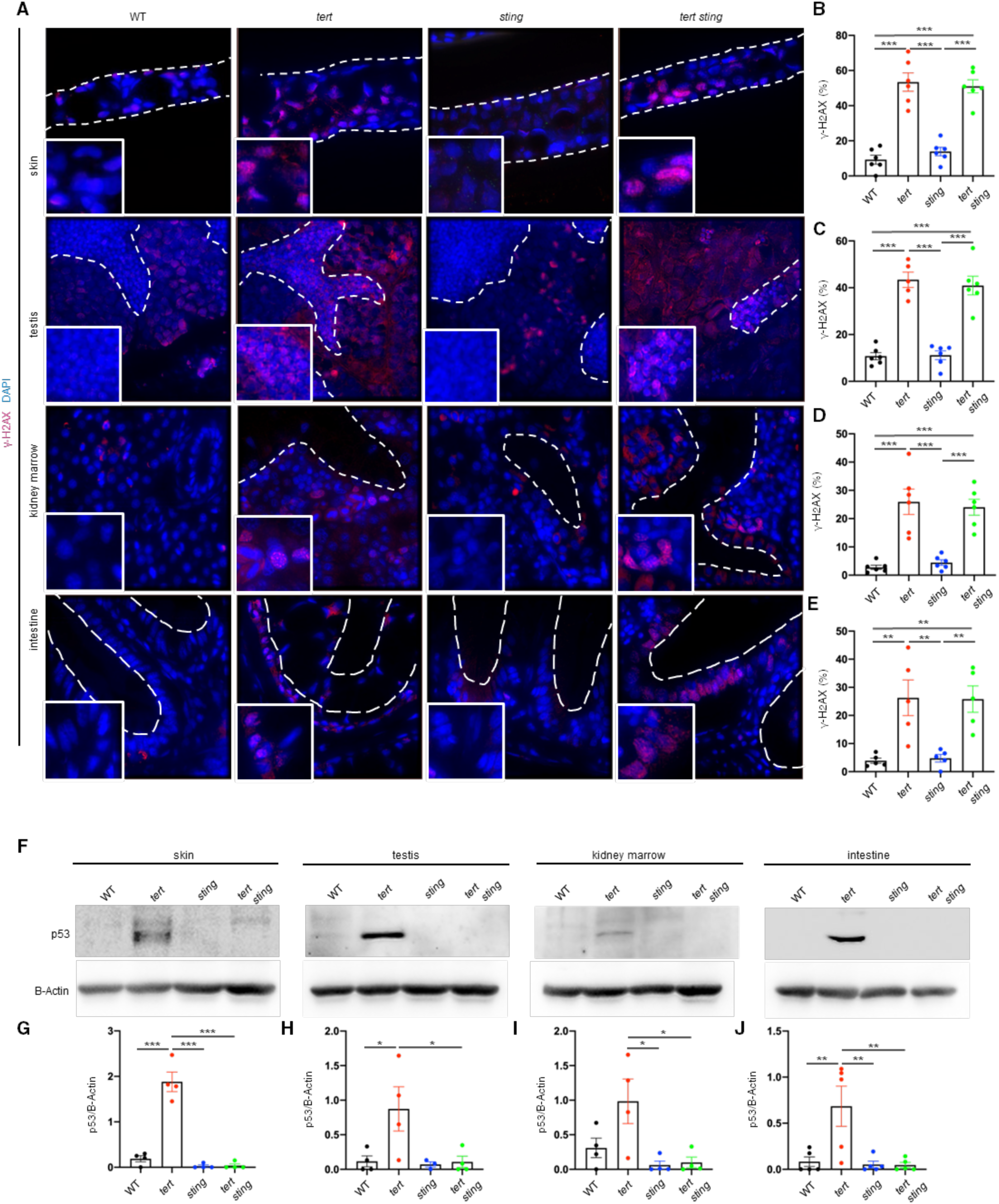
cGAS-STING pathway inactivation attenuates DNA damage response. **a**, representative immunofluorescence images of DNA damage. **b**, quantification of DNA damage in skin (n_WT_= 6, n*_tert-/-_*=6, n*_sting-/-_*=6, *_tert-/-sting-/-_*=6; WT vs tert-/- p< 0.001, sting-/- vs tert-/- p< 0.001, *WT* vs *tert-/- sting-/- p< 0.001, sting-/-* vs *tert-/- sting-/- p< 0.001*). **c**, quantification of DNA damage in testis (n_WT_= 6, n*_tert-/-_*=6, n*_sting-/-_*=6, *_tert-/-sting-/-_* =6, N= 5-6, WT vs *tert-/- p< 0.001, sting-/*v*-*s *tert-/- p< 0.001, WT vs tert-/- sting-/- p< 0.001, sting-/- vs tert-/- sting-/- p< 0.001*), **d**, quantification of DNA damage in kidney marrow (n_WT_= 6, n*_tert-/-_*=6, n*_sting-/-_* =6, *_tert-/-sting-/-_* =6; WT vs *tert-/- p< 0.001, sting-/*v*-*s *tert-/- p< 0.001, WT vs tert-/- sting-/- p< 0.001, sting-/- vs tert-/- sting-/- p<0.001*). **e**, quantification of DNA damage in intestine (n_WT_= 5, n*_tert-/-_*=5, n*_sting-/-_*=5, *_tert-/-sting-/_*=*_-_* 5; WT vs *tert-/- p=0.006, sting-/-* vs *tert-/- p=0.008, WT vs tert-/- sting-/- p= 0.007, sting-/- vs tert-/- sting-/- p<0.00*)*1*. **f**, representative western blot images of p53. **g**, quantification of p53 levels in the skin (n_WT_= 4, n*_tert-/-_*=4, n*_sting-/-_* =4, *_tert-/-sting-/-_*=4, WT vs *tert-/- p< 0.001, sting-/-*vs *tert-/- p< 0.001, tert-/-*vs *tert-/- sting-/- p< 0.001*). **h**, quantification of p53 levels in the testis (n_WT_= 4, n*_tert-/-_*=4, n*_sting-/-_*=4, *_tert-/-sting-/-_*=4, WT vs *tert-/- p=0.050, tert-/-* vs *tert-/- sting-/- p=0.048*). **i**, quantification of p53 levels in the kidney marrow (n_WT_= 4, n*_tert-/-_*=4, n*_sting-/-_*=4, *_tert-/- sting-/-_*=4, *sting-/-*vs *tert-/- p= 0.017, tert-/-*vs *tert-/- sting-/- p= 0.022*). **j**, quantification of p53 levels in the intestine (n_WT_= 5, n*_tert-/-_*=5, n*_sting-/-_*=5, *_tert-/-sting-/-_*=5, WT vs *tert-/- p<0.001, sting-/-*vs *tert-/- p=0.006, tert-/-*vs *tert-/- sting-/- p=0.006*). Data are presented as the mean ± s.e.m.; *p<0.05; **p<0.01, ***p<0.001, using a one-way ANOVA and post hoc Tukey test.

Increased γ-H2AX levels in *tert-/-* zebrafish is accompanied by elevation in p53 protein levels^8,21^. *tert-/-* zebrafish exhibited an increase of 5-fold (Figure 2F,G) in the skin and testis (Figure 2F,H), 3-fold in the kidney marrow (Figure 2F,I) and 6-fold in the intestine (Figure 2F,J) when compared to WT and *sting-/-* mutants (Figure 2E-H). Surprisingly, p53 levels in the *tert-/- sting-/-* mutants were similar to the WT and *sting-/-* siblings (Figure 2F-J). Our results indicate that, despite similar levels of phosphorylated H2AX, cGAS-STING is required for elevated p53 levels in response to telomere shortening. This is in agreement with studies showing that p53 expression and stability are regulated by IFNs^24,25^.

### cGAS-STING is required for senescence and SASP caused by telomere shortening

Given that p53 was not elevated in aging *tert-/- sting-/-* zebrafish, despite the shorter telomere length, we decided to assess the remaining phenotypes linked with *tert-/-* premature aging. Previous studies in human cells showed that cGAS-STING is required for cell senescence^10–12^. Using 9-months-old zebrafish, we studied cell senescence using the SA-Beta-Galactosidase (SA-B-gal) assay in addition to expression of *cdkn2a/b* (p15/16) and *cdkn1a* (p21) by RT-qPCR. As previously observed ^5,6,8,21^, proliferative tissues of *tert-/-* mutants were strongly stained with SA- B-gal, but not those of WT or *sting-/-* siblings (Figure 3A). Consistently, we confirmed an elevated expression of *cdkn2a/b* and *cdkn1a* senescence markers in *tert-/-* mutants (Figure 3B-D). Senescence in *tert-/-* was accompanied by expression of SASP-related genes, namely *il1b*, *tgfb1b* and *mmp15a*. As expected, none of these genes were elevated in WT or *sting*-/- siblings (Figure 3B-D).

**Figure 3:**
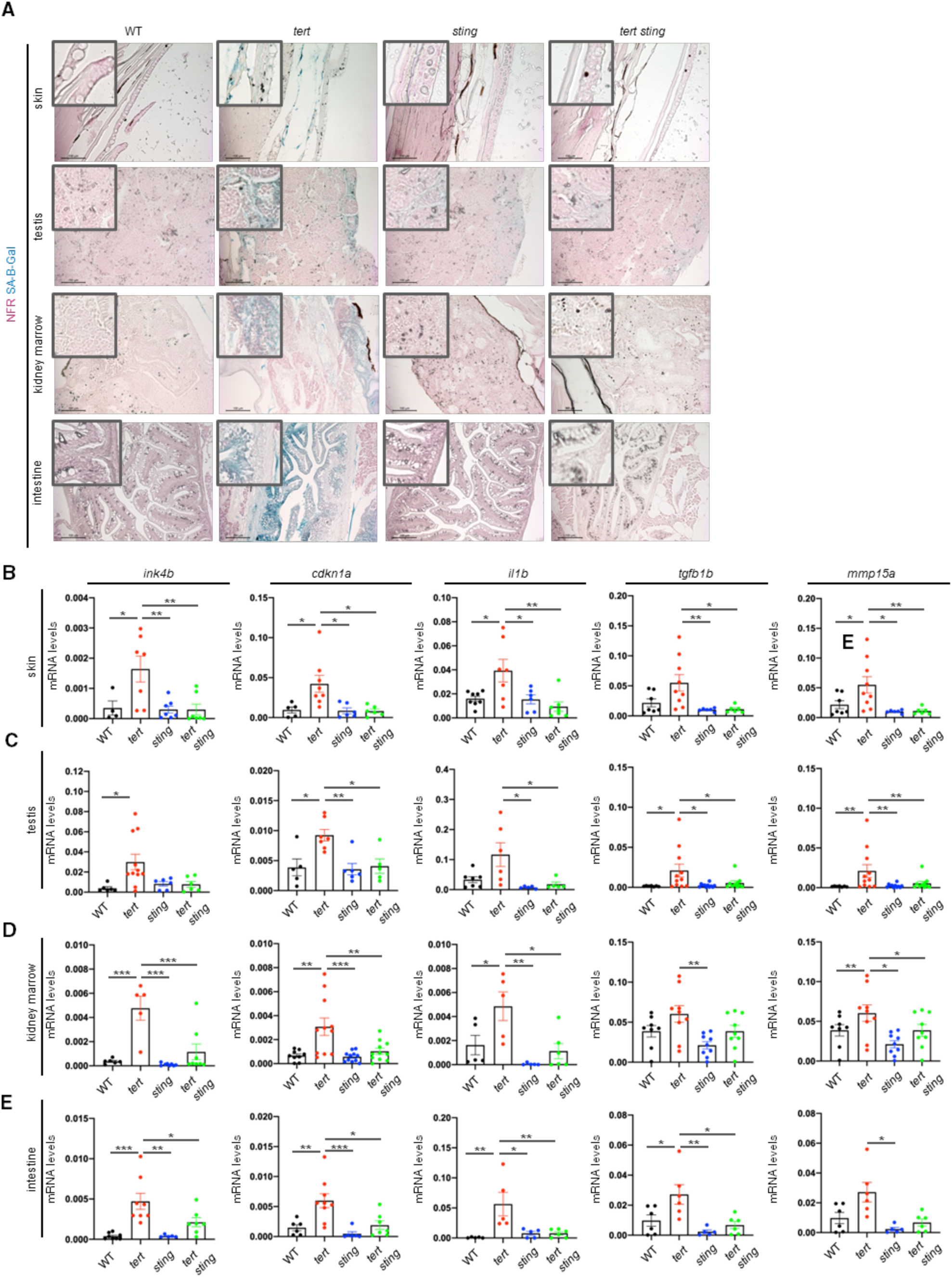
SASP induced by short telomeres are controlled by cGAS-STING pathway. **a**, representative images of senescence associated beta galactosidase staining in skin, testis, kidney marrow and intestine (n_WT_= 3, n*_tert-/-_*=3, n*_sting-/-_* =3, *_tert-/-sting-/_*=*_-_* 3). **b**, RT-qPCR analysis of inflammatory markers and SASP factors in the skin (n_WT_= 4-8, n*_tert-/-_*=5-8, n*_sting-/-_*=6-7, *_tert-/-sting-/-_*=5-7; *cdkn2a/b:* WT vs *tert-/- p=0.0001, sting-/-*vs *tert-/- p<0.0001, tert-/-*vs *tert-/- sting -/- p=0.004; cdnk1a:*WT vs *tert-/- p=0.025, sting-/-*vs *tert-/- p=0.014, tert-/-* vs *tert-/- sting-/- p=0.013; il1b*: WT vs *tert-/- p=0.023, sting-/-*vs *tert-/- p=0.031, tert-/-*vs *tert-/- sting-/- p=0.004,; tgf1b: sting-/-*vs *tert-/- p=0.014, tert-/-*vs *tert-/- sting-/- p=0.017; mmp15a: sting-/*v*-* s *tert-/- p=0.011, tert-/-*vs *tert-/- sting-/- p=0.009*). **c**, RT-qPCR analysis of inflammatory markers and SASP factors in the testis (n_WT_= 4-7, n*_tert-/-_*=6-11, n*_sting-/-_* =5-7, *_tert-/-sting-/_*=*_-_* 5-8; *cdkn2a/b:* WT vs *tert-/- p=0.0234; cdnk1a:*WT vs *tert-/- p=0.013, tert-/-*vs *tert-/- sting-/- p=*0.018*; il1b: tert-/-*vs *tert-/- sting-/- p=0.030; tgf1b:*WT vs *tert-/- p=0.029, sting-/-* vs *tert-/- p=0.020, tert-/-*vs *tert-/- sting-/- p=0.043; mmp15a:*WT vs *tert-/- p=0.004, sting-/-* vs *tert-/- p=0.016, tert-/-*vs *tert-/- sting-/- p=0.028*). **d**, RT-qPCR analysis of inflammatory markers and SASP factors in the kidney marrow (n_WT_= 5-9, n*_tert-/-_*=5-11, n*_sting-/-_*=6-11, *_tert-/-sting-/-_*=6-10; *cdkn2a/b:* WT vs *tert-/- p=0.035, sting-/-* vs *tert-/- p=0.009, tert-/-*vs *tert-/- sting-/- p<0.001; cdnk1a:*WT vs *tert-/- p=0.002, sting-/-*vs *tert-/- p=0.001, tert-/-*vs *tert-/- sting-/- p=0.005; il1b*: WT vs *tert-/- p=0.047, sting-/-*vs *tert-/- p=0.003, tert-/-* vs *tert-/- sting-/- p=0.014; tgf1b: sting-/-*vs *tert-/- p=0.005; mmp15a:*WT vs *tert-/- p=0.009, sting-/-* vs *tert-/- p=0.011, tert-/-*vs *tert-/- sting-/- p=0.008*). **e**, RT-qPCR analysis of inflammatory markers and SASP factors in the intestine (n_WT_= 5-7, n*_tert-/-_*=5-8, n*_sting-/-_*=5-6, *_tert-/-sting-/-_*=5-7; *cdkn2a/b:* WT vs *tert-/- p<0.001, sting-/-*vs *tert-/- p=0.001, tert-/-*vs *tert-/- sting-/- p=0.038; cdnk1a:*WT vs *tert-/- p=0.007, sting-/-*vs *tert-/- p<0.001, tert-/-* vs *tert-/- sting-/- p=0.011; il1b*: WT vs *tert-/- p=0.004, sting-/-*vs *tert-/- p=0.012, tert-/-*vs *tert-/- sting-/- p=0.009; tgf1b:* WT vs *tert-/- p=0.034, sting-/-*vs *tert-/- p=0.003; mmp15a:*WT vs *tert-/-* NS *p=0.060, sting-/-* vs *tert-/- p=0.021, tert-/-*vs *tert-/- sting-/- p=0.070*). All data are presented as the mean ± s.e.m.; *p<0.05; **p<0.01, ***p<0.001, using a one-way ANOVA and post hoc Tukey test

With lower levels of p53, cell senescence was also reduced in *tert-/- sting-/-* tissues, as observed by low SA-Beta-gal and expression of *cdkn2a/b* and *cdkn1a*. In absence of senescence, expression of SASP factors were also reduced in *tert-/- sting-/-* mutants to the ones observed in WT (Figure 3B-D). Thus, in agreement with previous *in vitro* studies, our results indicate that cGAS-STING is required for senescence and SASP of proliferative tissues *in vivo*.

### cGAS-STING controls cell proliferation and tissue integrity of *tert-/-* zebrafish

Replicative cell senescence is a barrier against cell proliferation in response to telomere shortening. We thus investigated if absence of senescence in *tert-/- sting-/-* mutants would result in increased cell proliferation. We examined proliferative tissues by immunofluorescence using antibodies against PCNA, a marker for cell proliferation. As previously reported^5,6,8^, cell proliferation of *tert-/-* zebrafish is significantly reduced in proliferative tissues when compared to WT and *sting-/-* siblings (Figure 4B-D). However, lower cell proliferation was recovered to WT levels in *tert-/- sting-/-* zebrafish (Figure 4B-D). Thus, with lower cell senescence and p53 levels, absence of cGAS-STING results in an increase in cell proliferation despite telomere shortening in telomerase-deficient zebrafish.

**Figure 4:**
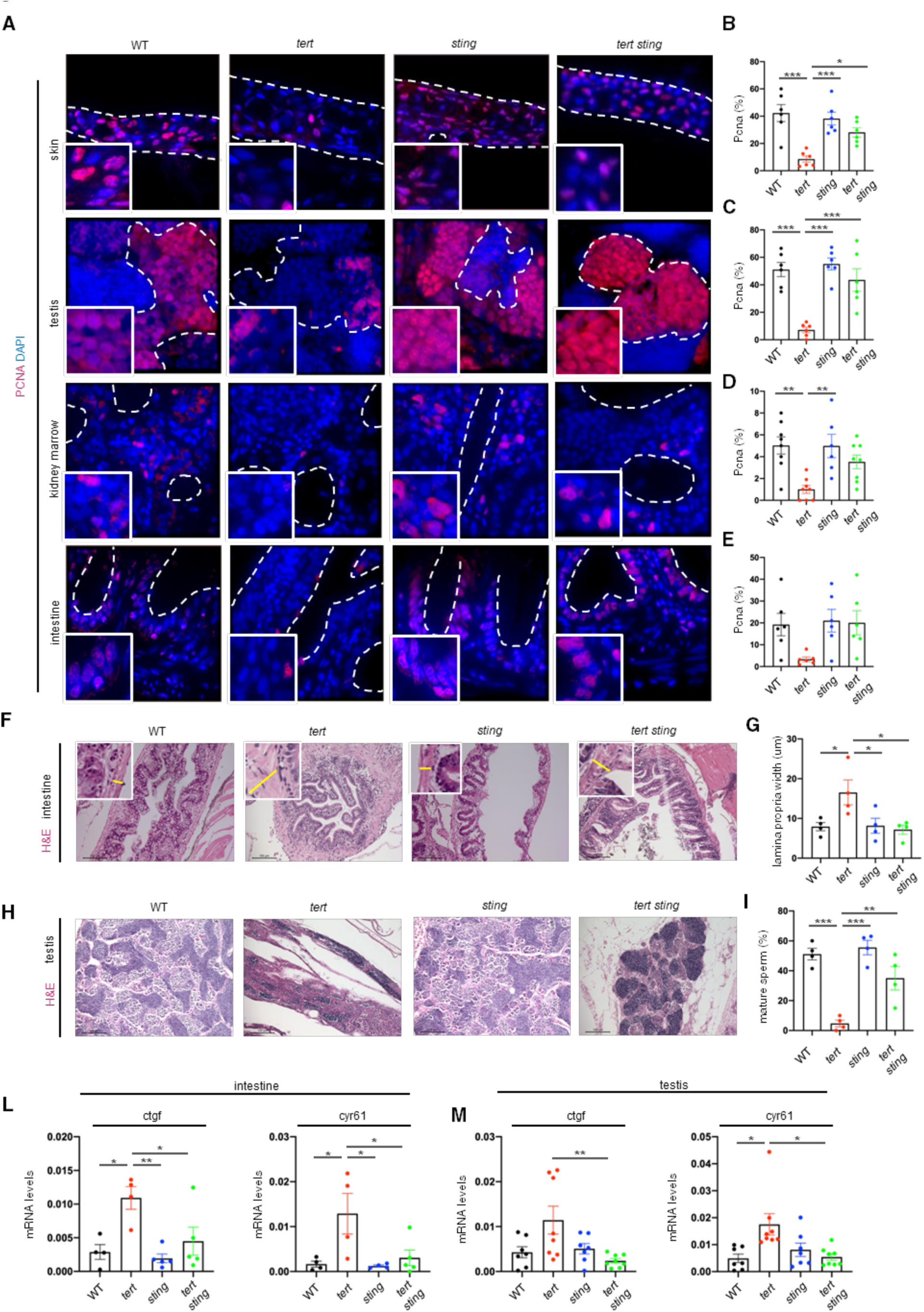
cGAS-STING pathway control proliferation in telomeric dysfunction. **a**, representative immunofluorescence images of proliferation and apoptosis. **b**, quantification of proliferation in the skin (n_WT_= 5, n*_tert-/-_*=6, n*_sting-/-_*=6, *_tert-/-sting-/-_*=6 WT vs *tert-/- p<0.001, sting-/-*vs *tert-/- p<0.001, tert-/-*vs *tert-/- sting-/- p=0.024*). **c**, quantification of proliferation in the testis (n_WT_= 6, n*_tert-/-_*=6, n*_sting-/-_*=6, *_tert-/-sting-/-_*=6, WT vs *tert-/- p<0.001, sting-/-*vs *tert-/- p<0.001, tert-/-*vs *tert-/- sting-/- p<0.001*). **d**, quantification of proliferation in the kidney marrow (n_WT_= 8, n*_tert-/-_*=8, n*_sting-/-_*=6, *_tert-/-sting-/-_*=8, WT vs *tert-/- p=0.002, sting-/-*vs *tert-/- p=0.004*). **e**, quantification of proliferation in the intestine (n_WT_= 6, n*_tert-/-_*=5, n*_sting-/-_*=6, *_tert-/-sting-/-_*=8). **f**, representative hematoxylin eosin staining of intestine, insets with yellow lines representative of lamina propria thickness **g**, quantification of lamina propria width (n_WT_= 4, n*_tert-/-_*=4, n*_sting-/-_*=4, *_tert-/-sting-/-_*=4, WT vs *tert-/- p=0.042, sting-/-* vs *tert-/- p=0.048*). **h**, representative hematoxylin eosin staining of testis. **i**, quantification of mature sperm area (n_WT_= 4, n*_tert-/-_*=4, n*_sting-/-_*=4, *_tert-/-sting-/-_*=4, WT vs *tert-/- p<0.002, sting-/-* vs *tert-/- p<0.001, tert-/-*vs *tert-/- sting-/- p= 0.006*). **l**, RT-qPCR analysis of YAP-TAZ pathway targets in the intestine (n_WT_= 4, n*_tert-/-_*=4, n*_sting-/-_*=5, *_tert-/-sting-/-_*=5, ctfg: WT vs *tert-/- p=0.016 sting-/-*vs *tert-/- p=0.005, tert-/-*vs *tert-/- sting-/- p=0.044; cyr61:* WT vs *tert-/- p=0.030, sting-/-*vs *tert-/- p=0,023, tert-/-*vs *tert-/- sting-/- p=0.046*). **m**, RT-qPCR analysis of YAP-TAZ pathway targets in the testis (n_WT_= 7, n*_tert-/-_*=8, n*_sting-/-_*=6-7, *_tert-/-sting-/-_*=7-8, cftg: *tert-/-* vs *tert-/- sting-/- p=0.008, cyr61: tert-/-*vs *tert-/- sting-/- p=0.011*). All data are presented as the mean ± s.e.m.; *p<0.05; **p<0.01, ***p<0.001, using a one-way ANOVA and post hoc Tukey test.

Previously, we showed that telomere shortening impacts tissue integrity and results in morphological defects of several tissues ^5,6,8,21^. Inflammation of the intestine causes an increase in thickness of the *lamina propria* in *tert-/-* zebrafish (mean of 17.5 micrometer, Figure 4F-G). However, width of the *lamina propria* in *tert-/- sting-/-* zebrafish was similar to the WT and *sting-/-* siblings (Figure 4G). Loss of gut tissue integrity activates the YAP-TAZ pathway in aging *tert-/-* zebrafish^8^. In agreement, we observed an increase of *ctfg* and *cyr6 1* levels, targets of YAP-TAZ pathway, (Figure 4L), in *tert-/-* zebrafish compared to WT and *sting*-/- siblings. However, expressions of YAP-TAZ target genes were reduced to the WT levels in the *tert-/- sting-/-* zebrafish (Figure 4L). Similarly, the remaining proliferative tissues also showed increased expression of *ctgf* and *cyr61* in *tert-/-* zebrafish that were absent in *tert-/- sting-/-* siblings (skin: 2.5-fold and 5-fold, respectively, kidney marrow: ∼10 fold, Figure 4M, Supplementary Figure 4B-D). Thus, our results suggest that tissue integrity of the intestine and other tissues caused by telomere shortening is rescued by an increase of cell proliferation upon inactivation of the cGAS-STING pathway.

### cGAS-STING causes premature aging of *tert-/-* zebrafish

Given the previous results at the cellular level, we investigated the consequences to the whole organism of absence cGAS-STING and Type I interferon in response to telomere shortening. We previously established male fertility and testicular atrophy as robust assays for aging zebrafish ^5,6,26^. To measure testicular atrophy, we quantified the mature sperm area in HE stained sections of whole testis (Figure 4H). Percentage of mature sperm area of 9-month-old *tert-/-* zebrafish was reduced to ∼5%, compared to ∼50% in WT and *sting-/-* siblings (Figure 4I). However, the percentage of mature sperm area increased to ∼40% in *tert-/- sting-/-* zebrafish (Figure 4I). Denoting the observed loss of tissue integrity in testis in *tert-/-* prematurely aged zebrafish, we saw increased expression of the YAP-TAZ pathway targets in *ctgf* and *cyr61* compared to WT and *sting-/-* siblings (Figure 4M). Consistent with our previous results, expression of *ctfg* and *cyr61* were reduced in *tert-/- sting-/-* to WT levels (Figure 4M).

We next asked if the rescue in the testis morphology of *tert-/- st i n*w*g*o*-*u*/*ld*-* restore fertility of aging *tert-/-* zebrafish. We previously reported that, from 6-month-old, *tert-/-* males become infertile paralleling the loss fertility observed in 18-month-old WT fish ^5^. As expected, 9- month-old *tert-/-* males were unable to produce fertilized eggs when crossed with young WT females (Figure 5A). Strikingly, *tert-/- sting-/-* males were still fertile even if slightly lower than the WT and *sting-/-* siblings (Figure 5A). Our data shows that inhibition of cGAS-STING and type I interferon response is sufficient to restore fertility to aging *tert-/-* zebrafish.

**Figure 5:**
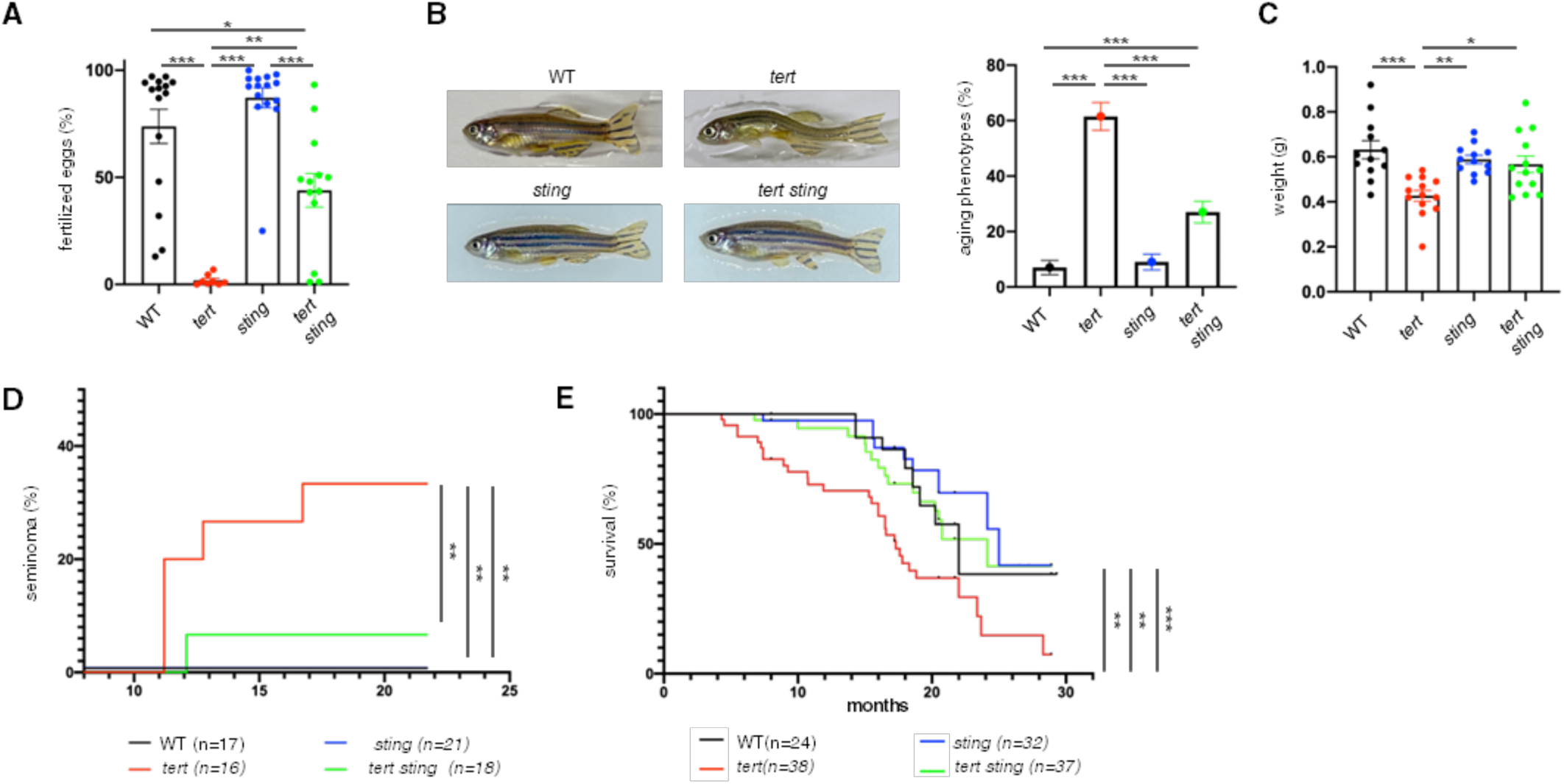
cGAS-STING pathway rescues premature aging phenotypes. **a**, quantification of male fertile capacity (n_WT_= 15, n*_tert-/-_*=10, n*_sting-/-_*=14, *_tert-/-sting-/_*=*_-_* 13), WT vs *tert-/- p<0.001, sting-/- vs tert-/- p<0.001,*WT vs *tert-/- sting-/- p=0.011, sting-/- vs tert-/- sting-/- p<0.001, tert-/*v*-*s *tert-/- sting-/- p=0.002)*. **b**, representative images of adult zebrafish and quantification of aging phenotypes scored as kyphosis and cachexia (n_WT_= 10, n*_tert-/-_*=12, n*_sting-/-_*=11, *_tert-/-sting-/-_*=12, WT vs *tert-/- p<0.001, sting-/- vs tert-/- p<0.001,*WT vs *tert-/- sting-/- p<0.001, sting-/-* vs *tert-/- sting-/- p=0.003, tert-/-*vs *tert-/- sting-/- p<0.001*). **c**, quantification of weight in adult zebrafish (n_WT_= 12, n*_tert-/-_*=13, n*_sting-/-_*=11, *_tert-/-sting-/-_*=13, WT vs *tert-/- p<0.001, sting-/- vs tert-/- p=0.004, tert-/-* vs *tert-/- sting-/- p=0.012*). **d**, quantification of seminoma (n_WT_= 17, n*_tert-/-_*=16, n*_sting-/-_*=21, *_tert-/-sting-/-_*=18, WT vs *tert-/- p=0.012, sting-/- vs tert-/- p=0.002, tert-/-*vs *tert-/- sting-/- p=0.027*). **e**, quantification of survival n_WT_= 24, n*_tert-/-_*=38, n*_sting-/-_*=32, *_tert-/-sting-/_*=*_-_* 37, WT vs *tert-/- p=0.008, sting-/- vs tert-/- p<0.001, tert-/-*vs *tert-/- sting-/- p=0.010*). Data are presented as the mean ± s.e.m. *p<0.05; **p<0.01, ***p<0.001, using a one-way ANOVA and post hoc Tukey test. Seminoma occurrence and survival data were analyzed using Log-rank tests, **p<0.01, ***p<0.001.

As other vertebrates, incidence of cancer increases with age in zebrafish ^5,26,27^. Similar to other age-associated phenotypes, the spontaneous cancer incidence is accelerated in younger ages in *tert-/-* zebrafish ^5,26^. We quantified the rate of spontaneous tumor formation, mostly seminomas, in male zebrafish. We observed that *tert-/-* zebrafish developed tumors from the age of 11-month-old and by the age of 17 months (Figure 5D). Strikingly, *tert-/- sting-/-* mutants developed macroscopic tumors around the age of 12 months but were restricted to 5% of the population (Figure 5D). Importantly, spontaneous cancer incidence of *tert-/- sting-/-* zebrafish was not statistically different from the WT and *sting-/-* siblings.

Other phenotypes of aging, such as kyphosis (abnormal curvature of the spine), caused by increased weakness of the spinal bones, and cachexia (excessive muscle wasting), caused by muscle tissue atrophy, are present in younger *tert-/-* mutants and only appear later in WT zebrafish ^5,6^. By the age of 15 months, 60% of *tert-/-* zebrafish showed aging phenotypes (Figure 5B) and weighed significantly less (Figure 5C) than WT and *sting-/-* siblings. However, the incidence of aging phenotypes ameliorated in *tert-/- sting-/-* mutants (Figure 5B) and weight was restored to WT levels (Figure 5C).

Finally, we compared the lifespan of *tert -*m*/*u*-*tants to their *tert-/- sting-/-* siblings. Whereas *tert-/-* mutants had a mean lifespan of 17 months-old, lifespan was extended to 24 months in *tert-/- sting-/-* mutants (Figure 5E). The mean lifespan of *tert-/- sting-/-* zebrafish was not statistically different from WT and *sting-/-* siblings (Figure 5F). Overall, our results show that by inhibiting cGAS-STING and, consequently, type I interferon, we observed an increase in lifespan of a prematurely aging vertebrate model by 41%. More importantly, we recover most age-associated phenotypes of aging *tert-/-* mutants, increasing their healthspan without the increase in cancer incidence.

## Discussion

Aging is accompanied by a wide range of physiological changes, such as chronic inflammation^28^. Age-associated inflammation in absence of overt infection has been termed inflammaging^29^. While the origins of inflammaging are mostly unclear, it is typically characterized by high levels of pro-inflammatory cytokines, chemokines, acute phase proteins, and soluble cytokine receptors in the serum ^29,30^. Inflammaging was shown to contribute to the development of age-associated diseases, such as neurodegenerative diseases, cardiovascular diseases and cancer ^28^. These are the main causes of morbidity and mortality in the elderly. Recent exciting new data revealed that DNA damage and inflammation are connected by the cGAS-STING pathway ^10,12,31^. Moreover, chemical inhibition of STING suppresses aging-associated inflammation and neurodegeneration^19^. Our study extends these observations by showing that activation of cGAS-STING pathway caused by telomere shortening is responsible for premature aging in zebrafish.

What triggers inflammaging upon telomere shortening? We documented several potential triggers for type I interferon responses in our work. We observed an increase of MN in *tert-/-* zebrafish. However, recent data has shown that, even though cGAS is recruited to MN, the presence of chromatin might not lead to elevated cGAMP and STING activation^32^. Further evidence from aging mice showed that mtDNA released from disrupted mitochondria was an important source for cGAS-STING activation^19^. In our previous work^21^, we showed that telomerase mutants have dysfunctional mitochondria with disrupted membranes providing a likely source for cytoplasmic mtDNA as an additional trigger. Moreover, mirroring what was previously observed in human fibroblasts^22,23^, we documented a tissue-specific elevation of TE expression, accompanied by increased expression of the RNA sensing pathway Rig-I, Mda5 and Mavs, that contribute and likely reinforce the activation type I interferon during aging. Finally, expression of the telomeric lncRNA TERRA could also contribute to the activation of cGAS-STING via the ZBP1/MAVS pathway, as reported for p53-deficient human cells undergoing crisis^23^. However, we were unable to detect a clear elevation of *mavs* expression in all tissues. A likely explanation may relate to *tert-/-* zebrafish being p53 proficient and, therefore, their comparatively longer telomeres may not express sufficient levels of TERRA to trigger its response.

Our work reveals that DDR and senescence triggered by short telomeres require an active cGAS-STING pathway. Even though telomere length and γ-H2AX levels were similar between *tert-/-* and *tert-/- sting-/-* zebrafish, p53 was only elevated in *tert-/-* mutants. Although a definite explanation is currently unavailable, we suggest potential leads for this observation. First, IFN-b signaling was shown to induce p53 transcription ^25^. Therefore, cGAS-STING may be required to promote p53 expression independently of canonical DDR. Second, the interferon-stimulated gene ISG15, an ubiquitin-like protein, is involved in p53 degradation by the 20S proteasome. ISG15 primarily targets misfolded p53 and deletion of ISG15 results in suppression of p53 activity and functions^24^. Thus, absence of cGAS-STING may lead to p53 destabilization. Third, cGAS-STING may sensitize cells to DNA damage by lowering the threshold for DDR activation. In absence of cGAS-STING, the ATM/ATR-p53 pathway may remain ineffective until genome instability is triggered by telomere-end fusions during crisis. In this scenario, phosphorylation of γ-H2AX could be achieved through parallel pathways, such as DNA-PKcs.

Absence of cGAS-STING in aging telomerase deficient zebrafish results in low p53 levels, reduced senescence and increased cell proliferation, thus rescuing damage imposed to proliferative tissues. This phenotype is also observed in *tp53-/- tert-/-* double mutant zebrafish^26^. Similar to late-generation telomerase deficient mice ^33^, lack of p53 leads to organismal rescue and increased fertility allowing for extra generations with ever-shorter telomere mice^34^. As first observed in tissue culture, upon telomere shortening, the first barrier to cell proliferation (M1) is imposed by p53/Rb and occurs when telomeres are long enough to allow for further cell divisions. ^35^. With loss of p53, the ensuing cell proliferation results in complete telomere deprotection, genome instability and cell death during crisis (M2)^31^. In this context, loss of cGAS-STING response is consistent with loss of p53 in aging *tert-/-* mutants. However, *tert-/- sting-/-* do not completely phenocopy *tert-/- tp53-/-* mutants. Loss of cGAS-STING does not cause an increase in spontaneous tumor incidence of either WT or *tert-/-* zebrafish. Zebrafish lacking p53, die prematurely primarily from increased soft tissue tumors^36^. In contrast, *tert-/- sting-/-* zebrafish lack elevated tumorigenesis characteristic of *tert-/- tp53-/-* mutants^26^. This may be attributed to the downstream consequences of cGAS-STING and type I interferon response. Chronic inflammation may be a key component of increased tumorigenesis in *tp53-/-* mutants. Lack of inflammatory responses may protect *tert-/- sting-/-* zebrafish from early tumorigenesis in face of increasing DNA damage.

We found that inhibiting cGAS-STING would restore tissue integrity and reduce expression of YAP-TAZ target genes in aging *tert-/-* mutants^8^. The YAP-TAZ pathway was recently shown to regulate cGAS-STING in stromal and contractile cells of aging mice^37^. Mechanotransduction by YAP-TAZ suppresses the activity of cGAS-STING, preventing senescence *in vivo* and tissue degeneration of prematurely aging mouse models ^37^. Our work now shows that cGAS-STING is also required for YAP-TAZ activity upon telomere shortening. This is likely to be the result of an indirect effect on tissue architecture. Restoring tissue integrity would reduce the activation of YAP-TAZ mechanosignaling and modifications to the extracellular matrix.

Loss of cGAS-STING and type I interferon improved the healthspan and the lifespan of aging *tert-/-* mutants. Telomere shortening does not occur simultaneously in all tissues in humans and zebrafish^5,38^. The gut of aging zebrafish presents early dysfunction and telomere shortening. We have recently shown that expressing telomerase specifically in the gut of *tert-/-* mutants prevents telomere shortening and gut dysfunction^8^. More importantly, maintaining telomere length in the gut also reverses remote organ dysfunction and longevity of the entire organism. Like *tert-/- sting-/-* zebrafish, gut-specific telomerase expression in *tert-/-* mutants reduces p53 levels and cell senescence in proliferative organs, namely testis and kidney marrow, despite short telomeres and increased levels of γ-H2AX. These lead to increased cell proliferation and restored tissue integrity. We propose that cGAS-STING and type I interferon responses initiated by telomere shortening in specific organs of aging individuals result in systemic chronic inflammation (inflammaging) deteriorating tissue integrity of remote organs by increasing DNA damage and reducing cell proliferation. Thus, inhibition of STING and chronic inflammation in organs primarily affected by telomere shortening, such as the gut and blood, would increase healthspan and lifespan. This provides a new approach for treatments of telomere biology disorders and to improve healthy aging.

## Materials and Methods

### Ethics statement

The zebrafish work was conducted according to local and international institutional guidelines and was approved in Portugal by the Ethics Committee of the Instituto Gulbenkian de Ciência and approved by the competent Portuguese authority (Direcção Geral de Alimentação e Veterinária; approval no. 0421/000/000/2015) and in France by the Animal Care Committee of the Institute for Research on Cancer and Aging, Nice, the regional (CIEPAL Côte d’Azur no. 697) and national (French Ministry of Research no. 27673-2020092817202619) authorities.

### Zebrafish lines and maintenance

Zebrafish were maintained in accordance with Institutional and National animal care protocols. To ensure telomere length comparisons and avoid the effects of haploinsufficiency of *tert+/-* heterozygous parental fish, we maintained double heterozygous stock lines (*tert*^AB/hu340^ *sting*^AB/sa35631^) as outcrosses to WT AB zebrafish. Experimental fish were obtained incrossing the stock fish. The overall characterization of these four genotypes was performed in F1 sibling animals at 9 months of age. Due to male sex bias in our crosses, that affected mostly *tert-/-* progeny, we were unable to obtain significant numbers of females for analysis and so all of our data except survival analysis are restricted to males.

### Telomere restriction fragment (TRF) analysis by Southern blot

Isolated tissues were first lysed at 50°C overnight in lysis buffer (Fermentas #K0512) supplemented with 1mg/ml Proteinase K (Sigma Aldrich) and RNase A (1:100 dilution, Sigma Aldrich). Genomic DNA was then extracted by equilibrated phenol-chloroform (Sigma Aldrich) and chloroform-isoamyl alcohol extraction (Sigma Aldrich). The same amount of gDNA was digested with RSAI and HINFI enzymes (NEB) for 12 h at 37°C. After digestion, samples were loaded on a 0.6% agarose gel, in 0.5% TBE buffer, and run on an electrophoresis apparatus (Bio-Rad). The electrophoresis conditions were 110 V for 15 h. Gels were then processed for Southern blotting using a 1.6 kb telomere probe, (TTAGGG)n, labeled with [alpha-32P]-dCTP.

### Fibroblast Derival

9-month-old fish zebrafish were sacrificed in 1g/L of MS-222 (Sigma Aldrich), and the skin was collected in PBS. After 3 washes in PBS + 2% antibiotics (Penicillin/Streptomycin, Gentamycin and Amphotericin b), the skin was dissociated for 5 minutes in Tryple (Gibco), cut into small pieces, and let to adhere O/N on coverslip coated by gelatin 2% in presence of few drops of FBS + 2% antibiotics. The day after, when the first fibroblasts were released from the skin, the well was filled with DMEM + 2% antibiotics. After 48h, fibroblasts were fixed and DAPI staining was performed.

### Western Blot

Age and sex matched adult zebrafish were sacrificed in 1g/L of MS-222 (Sigma Aldrich) and collected tissues (skin, testis, kidney marrow and intestine) were immediately snap frozen liquid nitrogen. Tissues were homogenized in RIPA buffer (sodium chloride 150 mM; Triton-X-100 1%, sodium deoxycholate 0.5%, SDS 0.1%, Tris 50 mM, pH=8.0), including complete protease and phosphatase inhibitor cocktails (Roche diagnostics) with a motor pestle on ice. Homogenized tissues were incubated for 30 minutes on ice and centrifuged at 4°C, 13.000 rpm for 10 min. Supernatant was collected and stored at-80°C until use.

For each sample, 50 μg of protein was loaded per well, separated on 10% SDS-PAGE gels and transferred to Nitrocellulose Membrane (BioRad #1620097). The membranes were blocked in 5% milk and then incubated with the primary antibody overnight at 4°C. Antibody complexes were visualized by enhanced chemiluminescence (ECL) after incubation with the appropriate HRP-conjugated secondary antibody. Antibodies concentrations: anti-p53 (1:1000, Anaspec, 55342), anti TBK1 (1:1000, CST, 3504), anti pTBK1 (1:1000, CST, 5483), anti IRF3 (1:1000, CST, 11904), anti pIRF3 (1:1000, CST, 29047).

### Real-time quantitative PCR

Age and sex matched adult zebrafish were sacrificed in 1g/L of MS-222 (Sigma Aldrich) and collected tissues (skin, testis, kidney marrow and intestine) were immediately snap frozen liquid nitrogen. RNA extractions were performed in TRIzol (Invitrogen) by mashing tissues with a motorized pestle in a 1.5 mL eppendorf tube. After incubation at room temperature (RT) for 10 min TRIzol, chloroform extractions were performed. Quality of RNA samples was assessed through BioAnalyzer (Agilent 2100). Retro-transcription into cDNA was performed using QuantiTect Reverse Transcription kit (Qiagen).

Quantitative PCR (qPCR) was performed using FastStart Universal SYBR Green Master mix (Roche) and an StepOne+ Real time PCR Detection System (Applied Biosystems). qPCRs were carried out in duplicate for each cDNA sample. Relative mRNA expression was normalized against *rps11* mRNA expression using the 2^-dCT^ method.

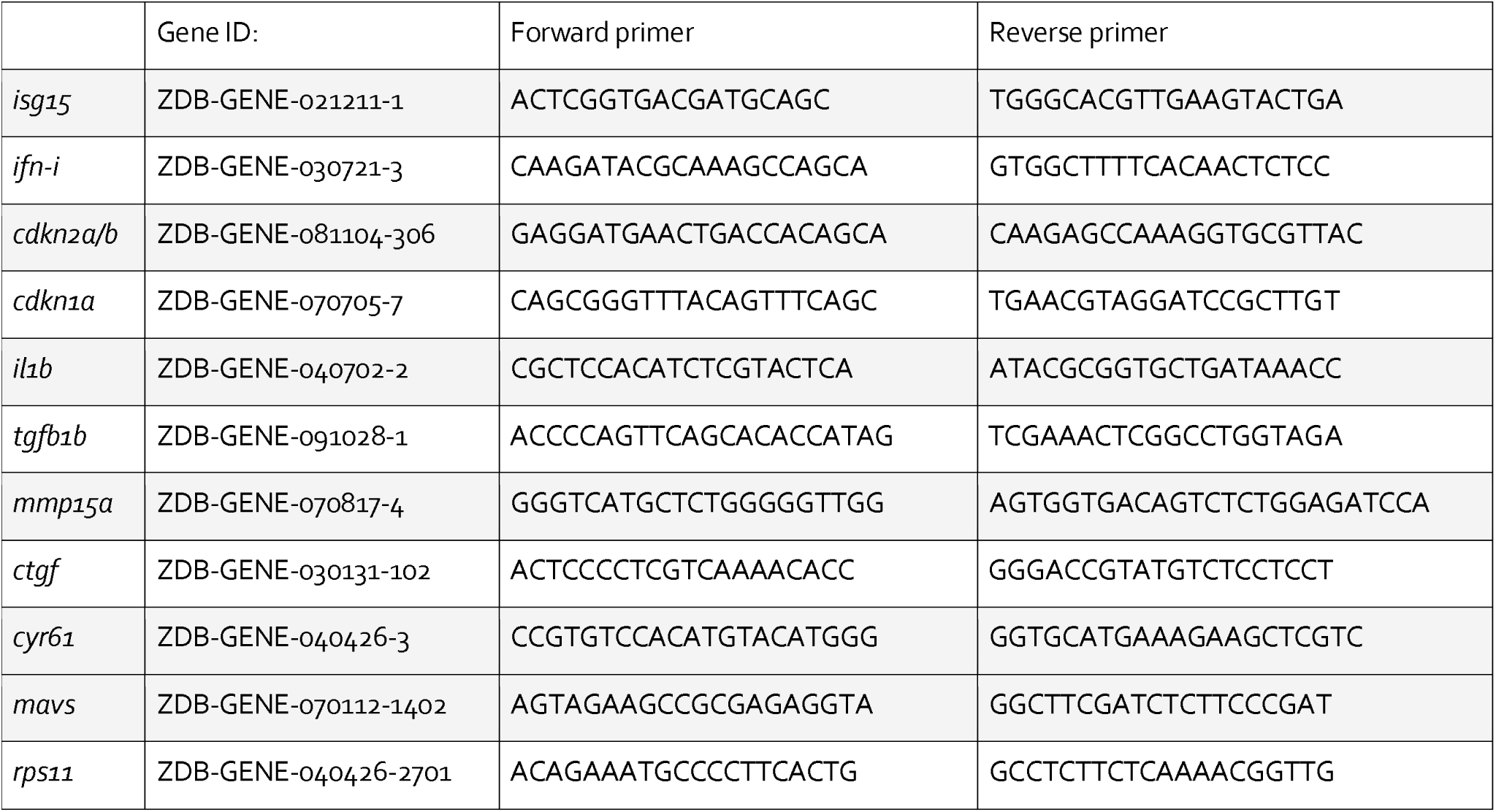

### Histology

Age and sex-matched adult zebrafish were sacrificed in 1g/L of MS-222 (Sigma Aldrich), fixed for 72 hours in 4% paraformaldehyde and decalcified in 0.5M EDTA for 48 hours at room temperature. Whole fish were then paraffin-embedded to perform five micrometer sagittal section slides. Slides were stained with hematoxylin (Sigma Aldrich) and eosin (Sigma Aldrich) for histopathological analysis. Microphotographs (N>=6 fish per genotype) were acquired in a Leica DM4000 B microscope coupled to a Leica DFC425 C camera (Leica Microsystems).

### Immunofluorescence

Deparaffinized and rehydrated slides were microwaved 20 min at 550W in citrate buffer (10 mM Sodium Citrate, pH 6.0) for antigen retrieval. Slides were washed twice in PBS for 5 minutes and blocked for 1 hour at RT in 0.5% Triton, 1% DMSO, 5% normal goat serum in PBS (blocking solution). Subsequently, slides were incubated overnight at 4°C with 1:50 dilution of primary antibody in blocking solution. The following primary antibodies were used: mouse monoclonal antibody against Proliferation Cell Nuclear Antigen (PCNA), sc56, (Santa Cruz), rabbit polyclonal Histone H2A.XS139ph (phospho Ser139) GTX127342 (Genetex). Following two PBS washes, overnight incubation at 4°C was performed 1:500 dilution of goat anti-rabbit secondary antibody AlexaFluor 488 (Invitrogen) and goat anti-mouse secondary antibody AlexaFluor 488 (Invitrogen). Finally, after DAPI staining (Sigma Aldrich), slides were mounted DAKO Fluorescence Mounting Medium (Sigma Aldrich).

Immunofluorescence images were acquired on Delta Vision Elite (GE Healthcare) using an OLYMPUS 60x/1.42 objective. For quantitative and comparative imaging, equivalent acquisition parameters were used. The percentage of positive nuclei was determined by counting a total of 150-1000 cells per slide depending on the tissue (N>= 6 zebrafish per genotype).

### Senescence Associated Beta Galactosidase Staining

Age and sex matched 9-month-old zebrafish were sacrificed in 1g/l of MS-222 (Sigma Aldrich, MO, USA), fixed in 4% paraformaldehyde in PBS for 72 hour at 4°C, washed 3 times for 1 hour in 1x PBS pH 7.4 and 1 hour in 1x PBS pH 6.0 at 4°C. Beta Galactosidase staining was performed for 24 hours at 37°C in 5mM Potassium Ferrocyanide, 5 mM Potassium Ferricyanide, 2 mM MgCl2 and 1 mg/mL X-Gal, in 1x PBS pH 6.0. After staining, fish were washed 3 times for 5 minutes in 1x PBS pH 7.0, and processed for decalcification and paraffin embedding. Paraffin blocks were sectioned sagittally five micrometers in thickness and co-stained with nuclear fast red (Sigma Aldrich)

### Fertility assays

In order to assess male fertility, 9-month-old males from the four different genotypes were separately housed overnight in external breeding tanks with a single young (3-6 months old) WT female. Breeding pairs were left to cross and lay eggs in the following morning and embryos were collected approximately 4 hours post fertilization (hpf) and allowed to develop at 28°C. Assessment of fertilized eggs and embryo viability was conducted between 4 and 6hpf. At least 12 independent crosses were conducted for each genotype to evaluate male fertility. Only successful breeding trials, defined as events in which a clutch of eggs was laid by a female, were scored.

### Fixation for histology and tumor evaluation

Fish were screened weekly for the presence of macroscopic tumors ^36^. Zebrafish were euthanized with 1g/L of MS-222 (Sigma, MO, USA), followed by fixation in 10% neutral buffered formalin for 72 hr and decalcified in 0.5M EDTA for 48 hr at room temperature. Samples were then paraffin-embedded to perform 5 mm sagittal section slides. Slides were stained with hematoxylin and eosin and assessed for the presence of tumors.

### RNA-seq analysis of TE families

Raw FASTQ read files obtained from the SRA (PRJNA937311) were aligned to the reference zebrafish genome (GRCz10/danRer10) using STAR v2.7.10a with the following parameters to retain multi-mapping reads: --outMultimapperOrder Random --outSAMmultNmax 1 -- outFilterMismatchNmax 3 --winAnchorMultimapNmax 100 --outFilterMultimapNmax 100 -- alignSJDBoverhangMin 1. Expression of transposable element families was quantified using TEtransctipts v2.2.393 with parameter –mode=multi to estimate transposable element abundances from multimapped alignments using the pre-generated GENCODE danRer10 TE GTF file from the Hammel lab FTP site (https://labshare.cshl.edu/shares/mhammelllab/www-data/TEtranscripts/TE_GTF). Counts were normalized to counts per million (CPM) and differential expression was assessed in R v3.1.4 (R Core Team, 2022) on the combined transposable element/gene counts using the edgeR package v3.34.189 exact test.

### Statistical Analysis

Graphs and statistical analyses were performed in GraphPad Prism8 software, using one-way ANOVA test Tukey’s post-correction or unpaired t-test. A critical value for significance of p<0.05 was used throughout the study. For survival analysis, Log-rank tests were performed using GraphPad Prism8 to determine statistical differences of survival curves.

## Data availability

All data generated or analyzed during this study are included in this published article and its Extended information files.

## Acknowledgments

We are grateful to our team members for fruitful discussions and advice. Especially, we thank Dr. Hervé Técher for critically reading the manuscript. This work was supported by the Université Co□te d’Azur - Académie 4 (Installation Grant: Action 2 - 2019) and Institut National du Cancer (INCa, PLBIO21-228). N.S. was supported by a PhD fellowship by La Ligue Contre le Cancer. E.T. and P.B. thank the German Research Foundation, grant nos. 322977937/GRK2344. We are grateful to the PEMAV fish facility, Imaging core facility (PICMI) and the Genomics facilities at the IRCAN supported by FEDER, Région Provence Alpes-Côte d’Azur, Conseil Départemental 06, ITMO Cancer Aviesan (plan cancer), Cancéropole Provence Alpes-Côte d’Azur, Gis Ibisa, CNRS and Inserm. The funders had no role in study design, data collection and analysis, decision to publish or preparation of the manuscript.

## Author contributions

N.S. performed most experiments and carried out data analyses. G.A. performed data analysis and revised the manuscript. B.L.-B. performed the skin fibroblast cell derivation and micronuclei quantifications. M.M. proposed the idea for the study and performed the initial analysis of the *sting-/- tert-/-* double mutants. RNA-seq analysis of transposon elements was performed by P.B. and supervised by E.T. N.S. and M.G.F. designed the experiments and wrote the manuscript. M.G.F. conceived the study, acquired funding and supervised the work.

