## Supplemental Figures for "cGAS-STING is responsible for aging of telomerase deficient zebrafish"

### Supplementary Figures and Legends

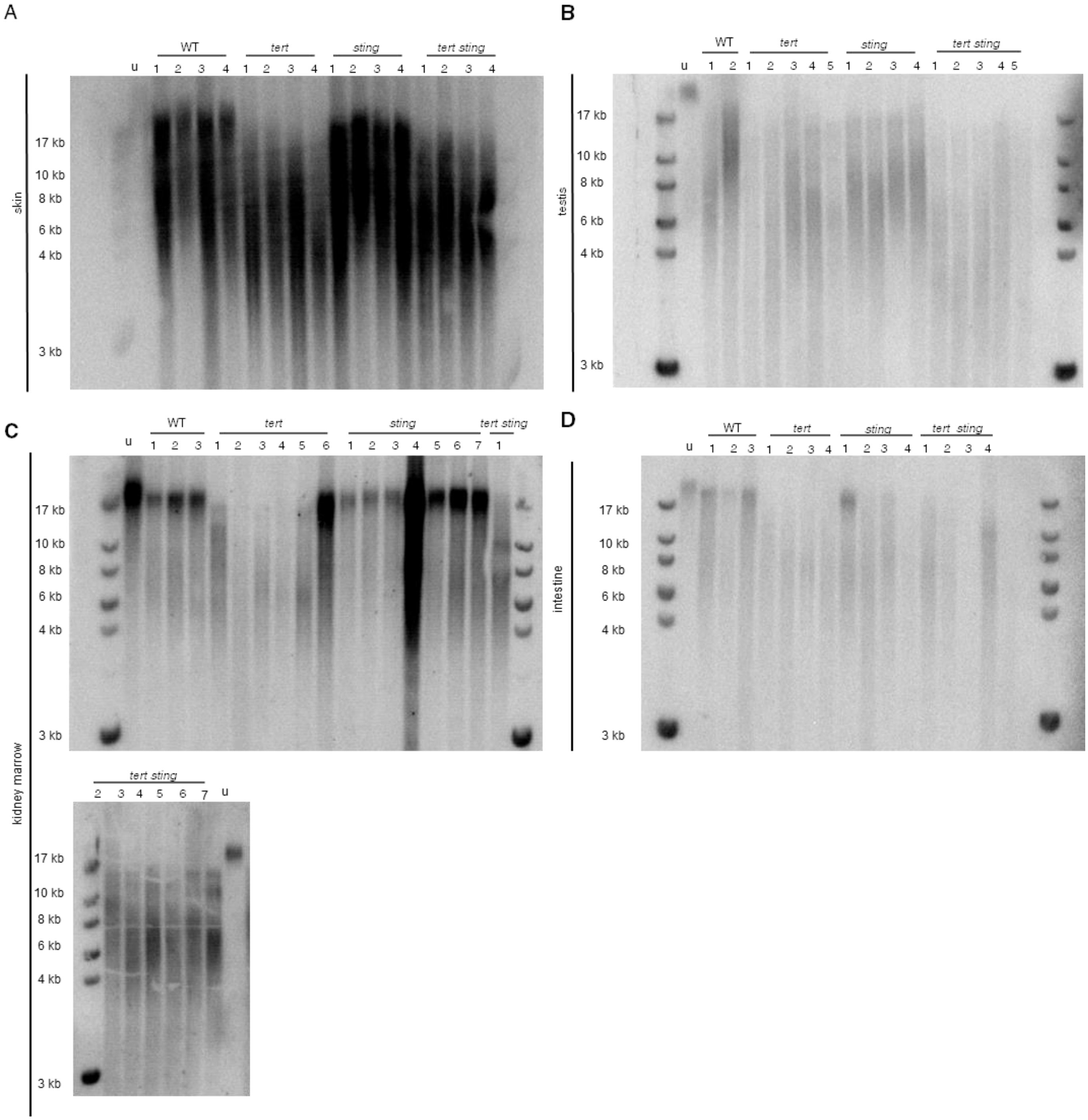


**Supplementary Figure 1: Southern blots for TRF assay** **a**, representative image for mean telomere length measured by TRF analysis in the skin ($n_{WT}$= 5, $n_{tert-/-}$=5, $n_{sting-/-}$=5, $_{tert-/-\ sting-/-}$=4). **b**, representative image for mean telomere length measured by TRF analysis in the testis ($n_{WT}$= 4, $n_{tert-/-}$=6, $n_{sting-/-}$=5, $_{tert-/-\ sting-/-}$=6). **c**, representative image for mean telomere length measured by TRF analysis in the kidney marrow ($n_{WT}$= 3, $n_{tert-/-}$=4, $n_{sting-/-}$=7, $_{tert-/-\ sting-/-}$=7) **d**, representative image for mean telomere length measured by TRF analysis in the intestine ($n_{WT}$= 6, $n_{tert-/-}$=6, $n_{sting-/-}$=6, $_{tert-/-\ sting-/-}$=6).

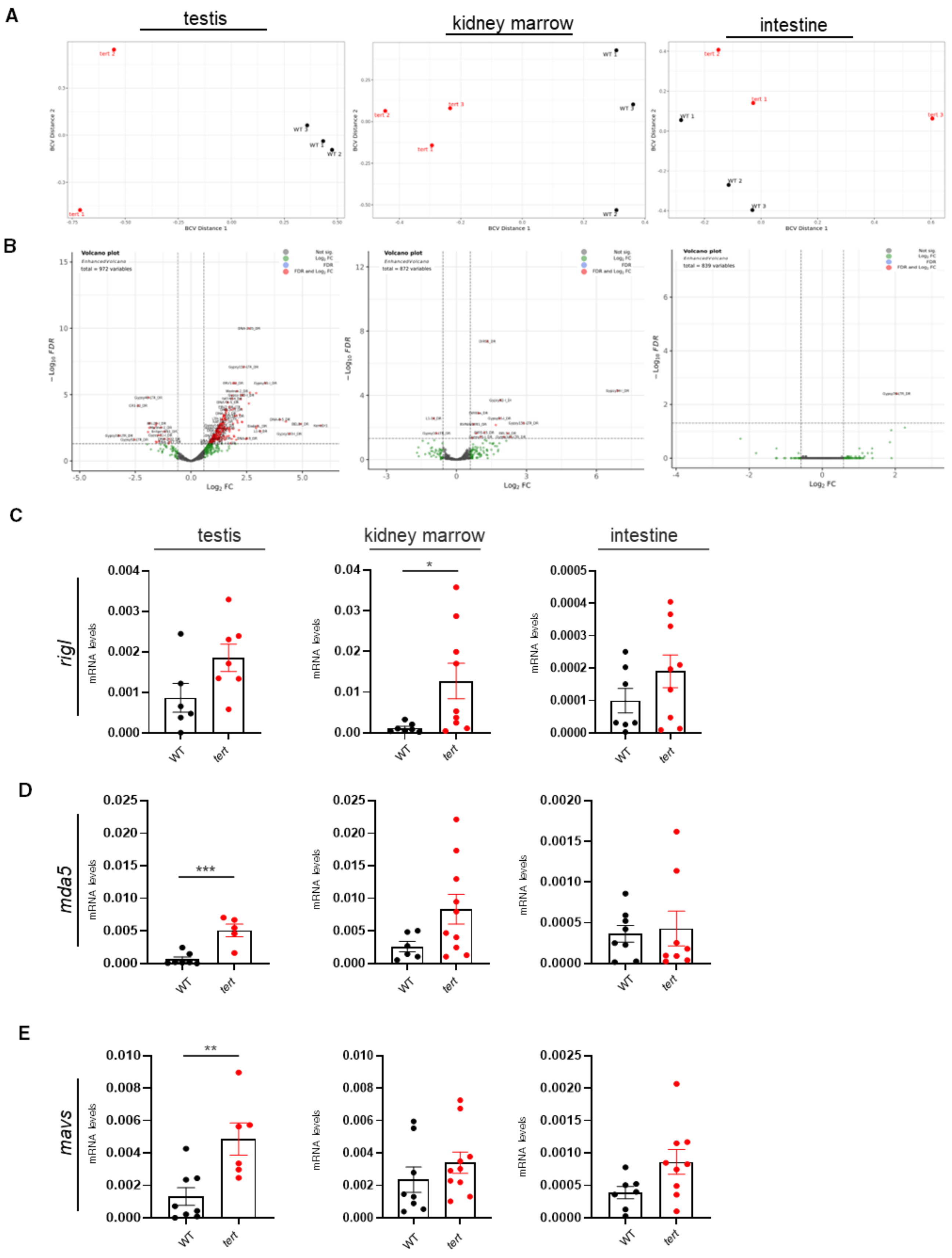


**Supplementary Figure 2: Expression levels of TEs and RNA sensor pathway. a,** Multidimensional scaling (MDS) plots of TE expression in WT and *tert* mutant fish. **b**, Vulcano plot of TE expressed in WT and *tert* mutant **c**, RT-PCR analysis of *rigI* gene expression in testis ($n_{WT}$= 8, $n_{tert\text{-}/\text{-}}$=8), in kidney marrow ($n_{WT}$= 8, $n_{tert\text{-}/\text{-}}$=9, p=0.038) and in intestine ($n_{WT}$= 7, $n_{tert\text{-}/\text{-}}$=9). **d**, RT-

PCR analysis of *mda5* gene expression in testis ($n_{WT}$= 8, $n_{tert-/-}$=6, p<0.001), in kidney marrow ($n_{WT}$= 6, $n_{tert-/-}$=10,) and in intestine ($n_{WT}$= 8, $n_{tert-/-}$=8,). **e**, RT-PCR analysis of *mavs* gene expression in testis ($n_{WT}$= 8, $n_{tert-/-}$=6, p=0.006), in kidney marrow ($n_{WT}$= 6, $n_{tert-/-}$=10,) and in intestine ($n_{WT}$= 8, $n_{tert-/-}$=8,). RNA expression data are presented as the mean ± s.e.m. *p<0.05; **p<0.01, ***p<0.001, using unpaired t-test.

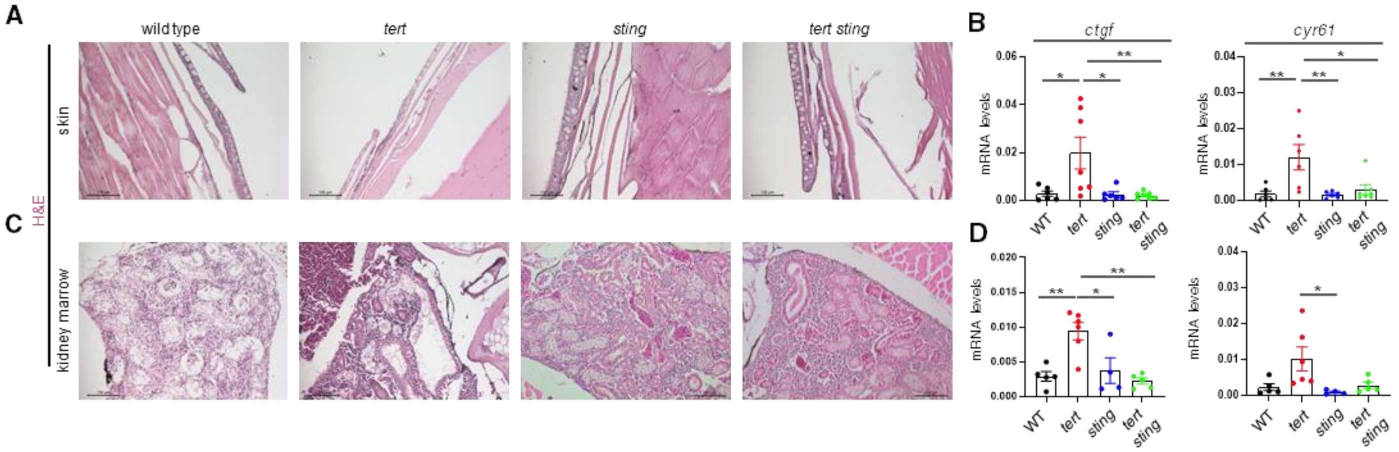


**Supplementary Figure 3: Histology and expression of YAP-TAZ targets. a** Representative hematoxylin eosin staining of skin ($n_{WT}$= 4, $n_{tert-/-}$=4, $n_{sting-/-}$=4, $_{tert-/-\ sting-/-}$=4). **b**, RT-qPCR analysis of YAP-TAZ pathway targets in the skin ($n_{WT}$= 6, $n_{tert-/-}$=6-7, $n_{sting-/-}$=6, $_{tert-/-\ sting-/-}$=7, *ctfg:* WT vs *tert-/- p=0.018, sting-/-* vs *tert-/- p=0.016, tert-/-* vs *tert-/- sting-/- p=0.010; cyr61:* WT vs *tert-/- p=0.007, sting-/-* vs *tert-/- p=0.006, tert-/-* vs *tert-/- sting-/- p=0.001*). **c**, representative hematoxylin eosin staining of kidney marrow ($n_{WT}$= 4, $n_{tert-/-}$=4, $n_{sting-/-}$=4, $_{tert-/-\ sting-/-}$=4). **d**, RT-qPCR analysis of YAP-TAZ pathway targets in the kidney marrow ($n_{WT}$= 5, $n_{tert-/-}$=6-7, $n_{sting-/-}$=4, $_{tert-/-\ sting-/-}$=5, *ctgf:* WT vs *tert-/- p=0.003, sting-/-* vs *tert-/- p=0.014, tert-/-* vs *tert-/- sting-/- p=0,001; cyr61: sting-/-* vs *tert-/- p=0.039*). Data are presented as the mean ± s.e.m. *p<0.05; **p<0.01, ***p<0.001, using a one-way ANOVA and post hoc Tukey test.
